# Social chemical communication determines recovery from L1 arrest via DAF-16 activation

**DOI:** 10.1101/2020.07.17.208066

**Authors:** Alejandro Mata-Cabana, Laura Gómez-Delgado, Francisco Javier Romero-Expósito, María Jesús Rodríguez-Palero, Marta Artal-Sanz, María Olmedo

## Abstract

In a population, chemical communication determines the response of animals to changing environmental conditions, what leads to an enhanced resistance against stressors. In response to starvation, the nematode *Caenorhabditis elegans* arrest post-embryonic development as L1 larva right after hatching. As arrested L1 larvae, *C. elegans* become more resistant to diverse stresses, allowing them to survive for several weeks expecting to encounter more favorable conditions. However, prolonged periods in L1 arrest lead to the accumulation of detrimental signs of aging, which ultimately provoke animal death. When arrested L1s feed, they undergo a recovery process to erase these harmful signs before resuming the developmental program. L1 arrested larvae secrete unidentified soluble compounds that improve survival to starvation. This protection is proportional to larval population density. Thus, animals arrested at high densities display an enhanced resistance to starvation. Here we show that this chemical communication also influences recovery after prolonged periods in L1 arrest. Animals at high density recovered faster than animals at low density. We found that the density effect on survival depends on the final effector of the insulin signaling pathway, the transcription factor DAF-16. Moreover, DAF-16 activation was higher at high density, consistent with a lower expression of the insulin-like peptide DAF-28 in the neurons. The improved recovery of animals after arrest at high density depended on soluble compounds present in the media of arrested L1s. In a try to find the nature of these compounds, we investigated the disaccharide trehalose as putative signaling molecule, since its production is enhanced during L1 arrest and it is able to activate DAF-16. We detected the presence of secreted trehalose in the medium of arrested L1 larvae at a low concentration. The addition of this concentration of trehalose to animals arrested at low density was enough to rescue DAF-28 production and DAF-16 activation to the levels of animals arrested at high density. However, despite activating DAF-16, trehalose was not capable of reversing survival and recovery phenotypes, suggesting the participation of additional signaling molecules. We finally identified GUR-3 as a possible trehalose receptor in *C. elegans*. With all, here we describe a molecular mechanism underlying social communication that allows *C. elegans* to maintain arrested L1 larvae ready to quickly recover as soon as they encounter nutrient sources.

## INTRODUCTION

Chemical communication between individuals of the same population determines the social behavior in response to environmental variations in a wide variety of species, from bacteria to mammals (Benabentos et al., 2009; Brückner and Parker, 2020; Coombes et al., 2018; Doyle and Meeks, 2018; Mukherjee and Bassler, 2019; Sokolowski, 2010; Taga and Bassler, 2003; Yan et al., 2020). Chemical communication in *C. elegans* has been mainly associated to ascaroside pheromones. These pheromones regulate a variety of social behaviors, including mating attraction, aggregation, foraging or avoidance (Artyukhin et al., 2013b; Butcher, 2017; Greene et al., 2016; Ludewig and Schroeder, 2013; MacOsko et al., 2009; Srinivasan et al., 2012, 2008). Ascarosides were firstly identified as social cues to regulate the entry in an alternative larval stage, *dauer*, in response to high-density population (Butcher et al., 2007; Golden and Riddle, 1982). In the form of *dauer* larvae, *C. elegans* can withstand harsh conditions for prolonged periods (reviewed in Hu, 2007). This exemplifies how social interactions can play a protective role in *C. elegans*, triggering a response to better resist unfavorable conditions. Animal aggregation in response to environmental stressors is another example of a protective social behavior, which allows *C. elegans* to improve survival until environmental conditions become more favorable (Boender et al., 2011; de Bono et al., 2002; Sokolowski, 2010).

As evolutionary strategy *C. elegans* larvae can arrest their development to overcome the detrimental effect of facing adverse conditions (Baugh et al., 2013). D*auer* arrest is induced in response to diverse environmental cues, such as high temperature, low food or high population density (Hu, 2007). However, post-embryonic development can also arrest at the first larval stage, L1, when animals do not encounter food right after egg hatching. When larvae enter the “L1 arrest” stage, cell proliferation stops and metabolism decreases. Arrested L1s can survive weeks without food and show increased resistance to stress. These worms must necessarily feed to resume development (reviewed in Baugh et al., 2013). Once these worms grow to adulthood they have normal lifespans, what was firstly interpreted as suspended aging during L1 arrest (Johnson et al., 1984). However, an aging process has been recently defined for these larvae, in which different detrimental signs of aging appear and accumulate during prolonged starvation (Roux et al., 2016). During a period of recovery after feeding these aging phenotypes are erased before the arrested L1s resume development (Roux et al., 2016). The length of the recovery time is directly proportional to the time in arrest and the grade of accumulation of signs of aging (Olmedo et al., 2020). However, long periods of L1 arrest impair the capacity of the larvae to recover (Roux et al., 2016).

The insulin/IGF-1 signaling pathway (IIS) is a major promoter of cell proliferation and growth in presence of food in *C. elegans*. In response to the secretion of insulin like peptides (ILP) by the sensory neurons, the insulin receptor DAF-2 gets activated and triggers a phosphorylation cascade. Low insulin signaling leads to the activation of the main final effector of the pathway, the transcription factor DAF-16, which migrates to the nucleus where it regulates the expression of multiple genes (reviewed in Murphy and Hu, 2013). DAF-16 is required for the larval survival to prolonged starvation (Baugh and Sternberg, 2006; Lee and Ashrafi, 2008; Muñoz and Riddle, 2003). DAF-16 also determines the aging process during L1 arrest and the recovery capacity of the larvae after starvation (Olmedo et al., 2020).

There are multiple factors affecting L1 survival to starvation, including environmental cues and social behavior (Artyukhin et al., 2015, 2013a; Baugh et al., 2013). Population density strongly influences L1 arrest survival (Artyukhin et al., 2013a). Larvae arrested at high densities display an enhanced resistance to starvation. This effect of larval density on survival depends on chemical communication between animals, in which the participation of sensory neurons is required. However, the identity of the signaling molecules involved remains unknown. It was determined that the nature of the signal was a small non-volatile molecule, but ascarosides were discarded as responsible of the density effect on L1 survival (Artyukhin et al., 2013a). *Caenorhabditis* species with density dependence on starvation survival also show L1 aggregation behavior on plates in the absence of food (Artyukhin et al., 2015, 2013a). Thus, both phenomena seem to be linked, and aggregation could be a way to locally increase population density, which enhances survival. However, the common thread between these two phenomena remains unidentified (Artyukhin et al., 2015).

Here we delve into the molecular mechanisms underlying the influence of population density and chemical communication on the capacity of arrested L1 larvae to resist and recover from long periods of arrest. We first show that high population density not only enhances survival to starvation but also enhances the capacity of larvae to recover from extended starvation. Both, survival and recovery from starvation are mediated by soluble compounds secreted to the medium. This means that social interaction not only improves the resistance of arrested L1s to prolonged starvation, but also keeps them ready to respond faster to a more favorable nutritional condition. We attribute a role for the IIS pathway in the social communication of the L1s. Animals arrested at high density present a lower insulin signaling that results in a higher activation of DAF-16, which is responsible for the enhanced recovery capacity after starvation. In an attempt to find communication molecules mediating the density effect, we investigated trehalose as a putative signal initiating the response. This disaccharide is present in the medium of arrested L1s at a low, density-dependent concentration. Our results show that trehalose mediates the communication between animals by reducing IIS signaling. The presence of trehalose in the medium leads to an inhibition in the production of the ILP DAF-28, and the activation of DAF-16. However, trehalose supplementation is not sufficient to reverse the shorter lifespan and longer recovery time of animals starved at low density, indicating that additional communication molecules are needed to completely explain the density effect on arrested L1 larvae. Finally, we also suggest GUR-3 as a possible trehalose receptor in *C. elegans* neurons.

## RESULTS

### Larval density during arrest affects recovery time

Prolonged periods of L1 arrest negatively influence starvation survival and the capacity of larvae to recover after refeeding (Olmedo et al., 2020; Roux et al., 2016). Larval density is an important factor affecting the survival of arrested L1 (Artyukhin et al., 2013a) and here we explored the possible influence of larval density during arrest in the recovery process. To do that, we measured recovery time of animals arrested at different densities by a quantitative method that precisely measures each stage of the post-embryonic development (Olmedo et al., 2015) (Fig. 1A). Thus, we arrested L1 larvae at 20, 2 and 0.2 L1/µl, during 1, 4, 8, or 14 days and then resumed development placing individual animals in wells of a 96 well plate with *E. coli*. This way, the effect of density during arrest is separated from a possible effect of density during recovery or its effect during developmental timing (Ludewig et al., 2017). When we looked at the effect of density during starvation on recovery time, we observed that animals arrested at lower densities required longer times to recover after extended starvation (Fig. 1B). After 8 days of starvation, the difference between the highest and the lowest density became statistically significant. Worms arrested at 0.2 L1/ µl needed, on average, 9.9 hours more to recover than those at 20 L1/ µl (33.1 h vs. 23.2 h) (Fig. 1B). Survival of animals arrested at 0.2 L1/ µl was dramatically impaired after 14 days on starvation. Although differences are not statistically significant, the trend to longer recovery times at lower densities is also observable in worms arrested at 2 L1/µl (Fig. 1B). Thus, worms at 2 L1/µl took, on average, 5.3 more hours to recover than those at high density (28.5 h vs. 23.2 h) after 8 days of arrest, and 13.8 hours more after 14 days of arrest (58.6 h vs. 44.8 h). Interestingly, developmental timing (M1 to M4) is not affected by larval density during arrest (Supp. Fig. 1A). The delay observed to complete the post-embryonic development (L1 to M4) at low densities can be completely explained by the longer recovery times (Supp. Fig. 1B). Therefore, as it happens for starvation survival, larval density also determines recovery after arrest.

**Figure 1.**
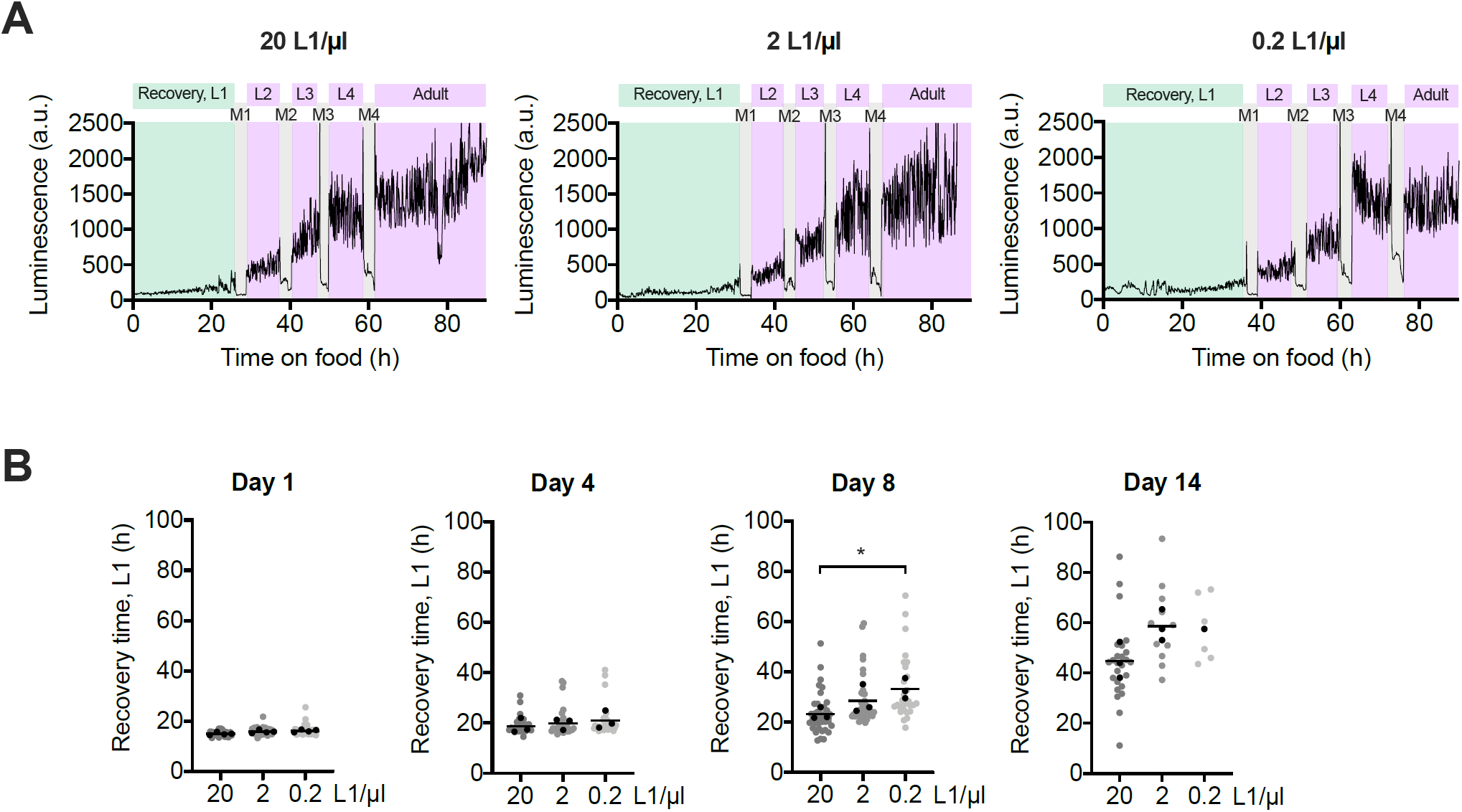
High population density enhances L1 recovery after prolonged starvation. (A) Representative luminometry profiles of development of animals arrested at 20, 2 and 0.2 L1/µl. Recovery time is colored in green and it is defined as the time from food addition to the first molt. The other larval stages and adulthood are colored in purple and the molts in grey. (B) Recovery times of animals arrested at densities of 20, 2 and 0.2 L1/µl after 1, 4, 8 and 14 days of starvation. All individual animals from three independent biological replicates are plotted. Black dots represent the average of each independent replicate and the black line indicates the mean of the entire experiment. Averages of the replicates were used for statistics (* p<0.05, Unpaired t-test). Supp. Fig. 1A and B respectively show the plots for developmental timing (M1 to M4) and the complete post-embryonic development (L1 to M4).

### DAF-16 mediates larval density effects on recovery from arrest

We recently showed that recovery time after extended arrest is regulated by Insulin/IGF-1-like signaling (IIS) in a DAF-16 dependent manner, being shorter in a *daf-2* mutant and longer in a *daf-16* mutant strain (Olmedo et al., 2020). At the same time, IIS also impacts survival of arrested L1s to prolonged starvation. Low insulin signaling results in a survival improvement, which is completely abolished by the loss of DAF-16 (Baugh and Sternberg, 2006; Lee and Ashrafi, 2008; Muñoz and Riddle, 2003). Arrested L1 larvae at low density show comparable phenotypes to *daf-16* mutant, higher mortality and a longer recovery time. Furthermore, mutants affected in the chemosensory proteins DAF-6, OSM-1 and TAX-4, which are necessary for the density-dependent effect (Artyukhin et al., 2013a), have an increased expression of *daf-28*, the gene encoding one of the *C. elegans* insulin-like peptides (ILPs) (Li et al., 2003). DAF-28 is an agonist ILP that promotes insulin signaling by binding and activating DAF-2 and, therefore, it represses DAF-16 activation (Kaplan et al., 2019; Li et al., 2003). Taking all this into consideration, we decided to address the question of DAF-16 as possible determinant of the effect of density on L1 survival. With this aim, we compared longevities of L1 larvae of wild-type and *daf-16* deletion strains arrested at different densities (Fig. 2A). To do that we transferred larvae from M9 buffer to NGM plates with food at the indicated times and determined the percentage of arrested L1s able to recover from arrest and complete development. Wild type larvae recovery was directly proportional to the density (Fig. 2A), and there was a progressive reduction in the half-life from the highest to the lowest density (Fig. 2B). Thus, the half-life of worms arrested at 2 L1/µl was around 7 days shorter than those arrested at 20 L1/µl, and larvae arrested at 0.2 L1/µl showed a half-life almost 6 days shorter than those at 2 L1/µl (Fig. 2B). With this experiment we also confirmed the impaired survival of the *daf-16* mutant L1 arrested larvae to prolonged starvation (Fig. 2B). At the highest density, the wild-type strain showed a half-life of around 23 days on average, and the half-life of the *daf-16* mutant was of around 7 days. Moreover, loss of DAF-16 completely abolished the density effect on L1 survival to starvation, since no differences are observed between the longevities of mutant larvae arrested at the three densities tested (Fig. 2A and B). In terms of longevity, the lowest density treatment applied to the wild type is not as severe as the loss of DAF-16, being the half-life of the mutant at high density in the range of 3 days shorter than the wild type at 0.2 L1/µl.

**Figure 2.**
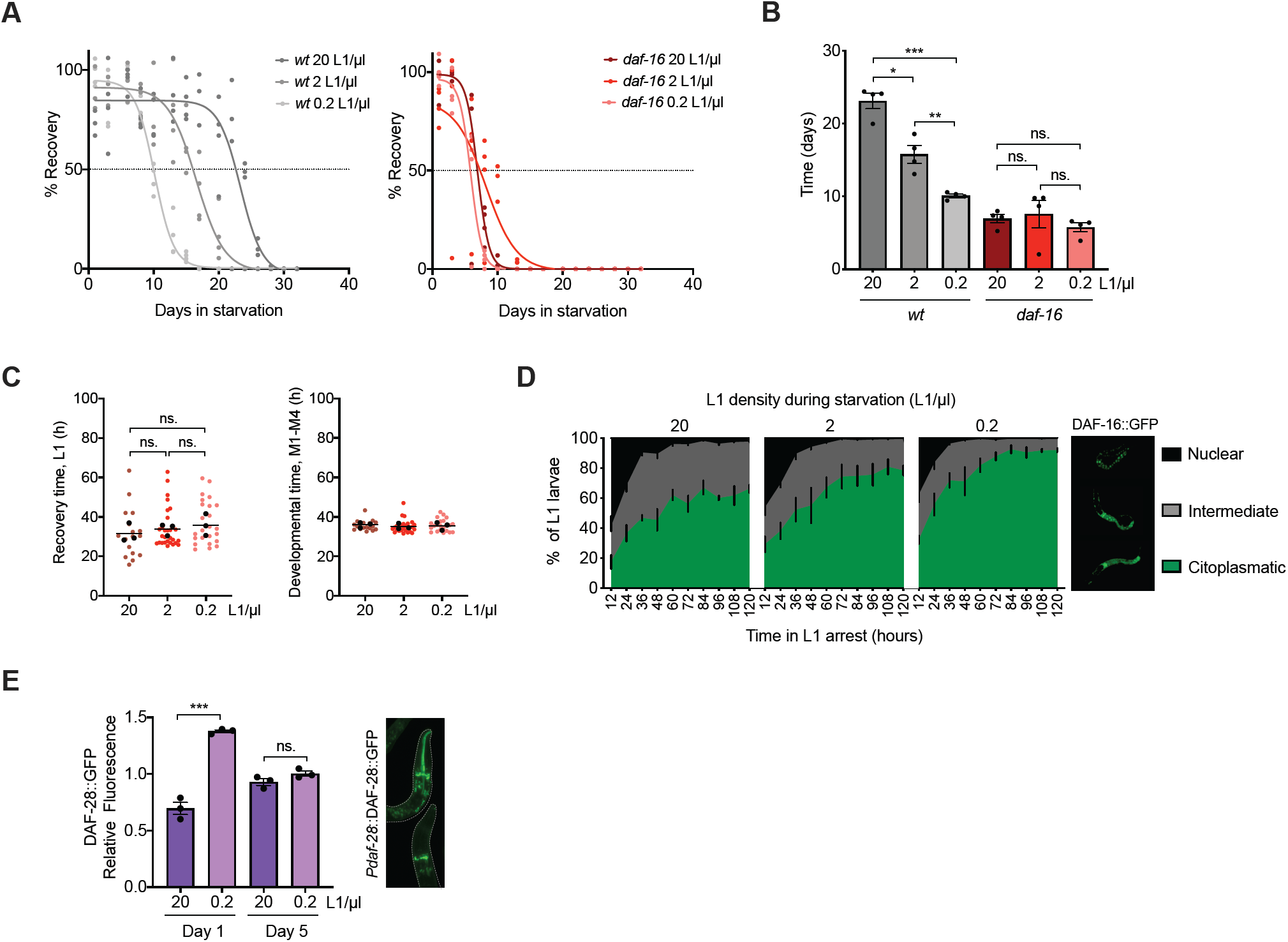
The density effect is mediated by DAF-16 activation. (A) Percentage of animals arrested at densities of 20, 2 and 0.2 L1/µl that recover after different periods of starvation. Survival curves for wild-type strain (left), and *daf-16* mutant (right). Each percentage calculated every 2-3 days is independently plotted as colored dots for four biological replicates. Curve fitting was performed using the data from the four replicates. The horizontal dotted line marks the 50% recovery, referred in the text as half-life. (B) Comparison of the half-lives in days of wild-type and *daf-16* strains arrested at 20, 2 and 0.2 L1/µl. Black dots represent the value for each independent replicate. The bars mark the mean of the entire experiment and error bars shows the SEM (* p<0.05, ** p<0.01, *** p<0.001, one-way ANOVA).(C) Recovery time (L1) and developmental time (M1-M4) of *daf-16* mutant arrested at 20, 2 and 0.2 L1/µl after 4 days of starvation. Each colored dot represents the data from a single animal analyzed from three independent biological replicates. Black dots represent the average of each independent replicate and the line marks the mean of the entire experiment. Averages of the replicates were used for statistics (one-way ANOVA). (D) Subcellular localization of DAF-16::GFP in L1 larvae arrested at densities of 20, 2 and 0.2 L1/µl for 1 to 5 days in starvation. Statistics are shown in Supp. Fig. 1C. (E) Quantification of the expression of DAF-28::GFP in the heads of L1 larvae arrested at 20 and 0.2 L1/µl for 1 or 5 days. Black dots represent the average of each independent replicate. The bars mark the mean of the entire experiment and error bars shows the SEM (*** p<0.001, Unpaired t-test).

The density effect on recovery time was observable in the wild type only after 8 days of arrest (Fig. 1B). At this time of arrest, the percentage of animals that recover is 83,6%. However, to assess this effect in the *daf-16* mutant we could not use such a long starvation period, since at this time arrested L1 larvae of this mutant barely survive. Instead, we analyzed recovery time in the *daf-16* mutant after 4 days of starvation, the rate of recovery is at 85 %, indicating a similar situation to the wild-type strain at day 8 (Fig. 2A). By doing this we found that there were no differences in recovery time between *daf-16* animals at the different densities tested after 4 days of arrest (Fig. 2C). At all densities, *daf-16* mutants needed a time to recover of about 33 hours, similar to the wild type arrested for 8 days at the lowest density tested. This result shows that the density effect on recovery time after starvation depends on DAF-16.

Due to the lack of density effect in the *daf-16* mutant we asked whether the density could be reflected in a different activation level of this transcription factor. To address this question, we used a reporter strain expressing DAF-16 fused to GFP, which allows the visualization of the subcellular localization of DAF-16. We quantified the percentage of animals containing the activated version of the protein. Doing this quantification every 12 hours for 5 days, we observed that the nuclear localization of DAF-16 was higher at high density from the beginning of the experiment and it was more sustained during all the period of starvation when compared to the lower densities tested. The level and speed of DAF-16 inactivation was inversely proportional to the population density (Fig. 2D, Supp. Fig. 1C). Therefore, this result suggests the presence of a signaling molecule in the medium that eventually triggers the activation of DAF-16, which exerts its protective role against starvation.

The lack of the density effect in *tax-4, osm-11* and *daf-6* mutants, which have increased DAF-28 (Artyukhin et al., 2013a), suggests that this ILP could be also involved in the effect of density. Since activation of IIS by DAF-28 leads to inactivation of DAF-16 (Li et al., 2003), this ILP seems a plausible candidate to mediate the density effect on DAF-16 activation. To confirm this mechanism, we analyzed the expression of DAF-28 using a fluorescent reporter, in which the peptide is fused to GFP and it is under the control of its endogenous promoter. We quantified the signal in the head of individual worms arrested at high density, 20 L1/µl, and low density, 0.2 L1/µl, after 1 and 5 days of arrest (Fig. 2E). We found a higher DAF-28 production in L1 larvae at low density after 1 day of arrest, which correlates with a lower activation of DAF-16 at this density. After 5 days of arrest the difference in DAF-28 production between densities disappeared. The signal measured at low density decreased from 1 to 5 days of arrest. However, the DAF-28::GFP signal at high density after 5 days of arrest is higher than after 1 day of arrest, what would suggest a partial loss of the IIS repression during prolonged starvation periods. This correlates with the gradual increase in the cytoplasmic localization of DAF-16 during the arrest. Nevertheless, the lower DAF-28 production after 5 days of starvation, when we find the lowest activation of DAF-16, reveals that other mechanisms different to the increase in DAF-28 production might be involved in DAF-16 inactivation during prolonged starvation.

### Secreted soluble compounds promote recovery and activate DAF-16 in arrested L1 larvae

As previously exposed, a signaling molecule of unidentified nature is responsible for the density effect on L1 starvation survival (Artyukhin et al., 2013a). When conditioned medium (CM), collected from L1 cultures arrested at high density, was used to arrest larvae at low densities, it rescued the survival impairment proper of low population density (Artyukhin et al., 2013a). With this precedent, we asked whether the high concentration of the unknown signaling molecule contained in the CM of larvae arrested at high density was also capable of reducing the recovery time of worms arrested at low density. We found that CM shortened the recovery time of L1 at low density to the level of animals arrested at high density for 8 days (Fig. 3A). Therefore, communication between animals through soluble compounds is important to improve survival to starvation and recovery from arrest after refeeding.

**Figure 3.**
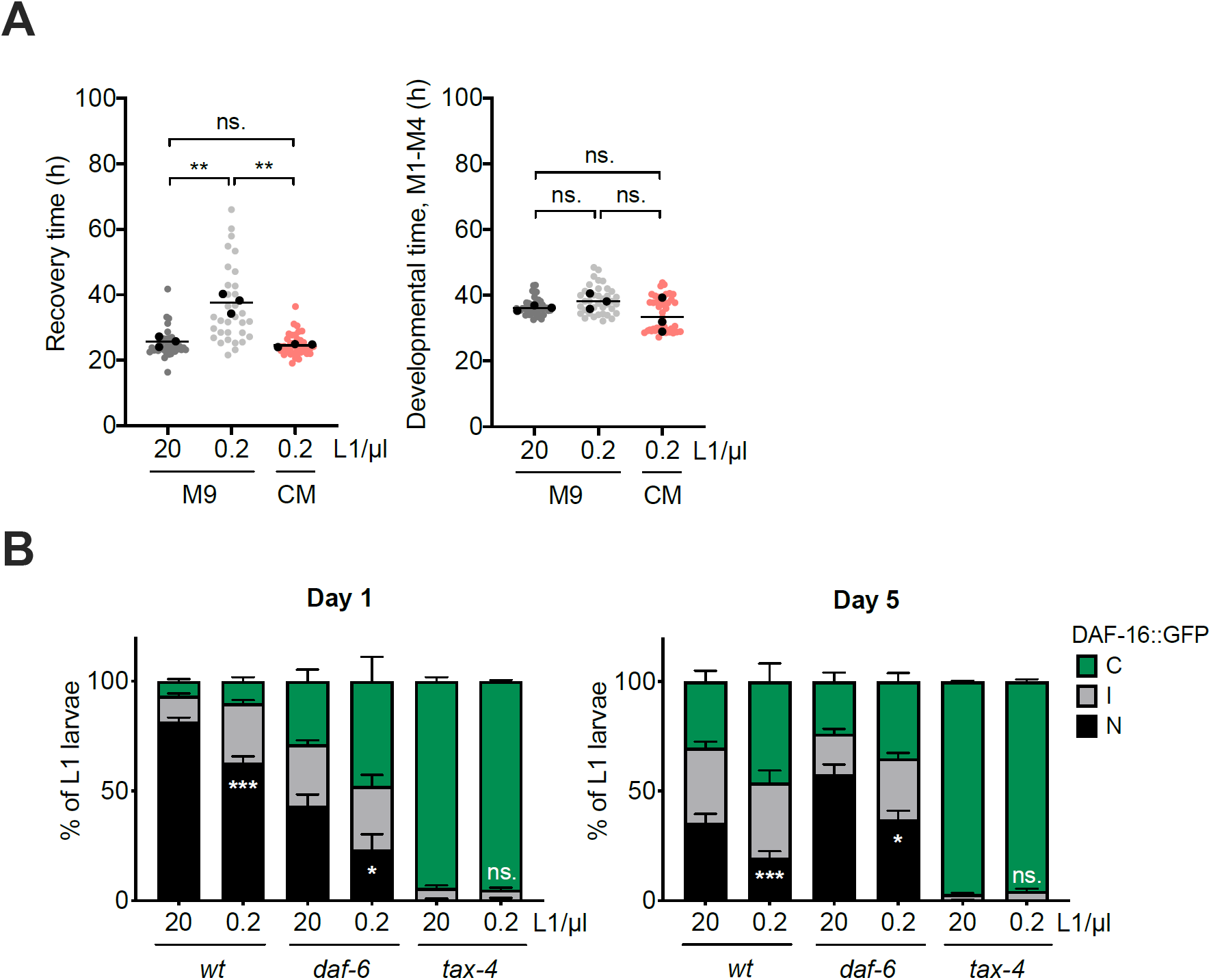
Chemical communication determines recovery and DAF-16 activation. (A) Comparison of recovery time (L1) and developmental time (M1-M4) after 8 days of starvation between animals arrested at 0.2 L1/µl in conditioned medium (CM) and animals arrested in M9 buffer at 20 and 0.2 L1/µl. Each colored dot represents the data from a single animal analyzed from three independent biological replicates. Black dots represent the average of each independent replicate and the line marks the mean of the entire experiment. Averages of the replicates were used for statistics (** p<0.01, Unpaired t-test). (B) Subcellular localization of DAF-16::GFP in L1 larvae arrested at 20 and 0.2 L1/µl for 1 and 5 days in *wt, daf-6* and *tax-4* mutant strains. The histograms show the means of, at least, three independent replicates and the error bars represent the SEM. Statistics for nuclear localization of DAF-16 are shown (* p<0.05, *** p<0.001, Unpaired t-test). Independent plots for the three localization categories are depicted in Supp. Fig. 2A.

We next analyzed the density dependence of DAF-16 activation in mutants of two of the chemosensory proteins, DAF-6 and TAX-4, described to be involved in the density effect (Artyukhin et al., 2013a). Both mutants, together with *osm-1* mutant, are defective in sensory neuron functioning and present an elevated *daf-28* expression level (Li et al., 2003). According to our results, DAF-16 showed a lower activation level in both mutants compared to the wild-type strain after 1 day of arrest (Fig. 3B, Supp. Fig. 2A). The effect was more dramatic in *tax-4* mutant, where the nuclear form of DAF-16 was inappreciable at both high and low density, what could explain the loss of density effect in L1 starvation survival. However, we could still observe an effect of density in the activation of DAF-16 in the *daf-6* mutant. Nevertheless, the higher DAF-16 nuclear localization at high density in this mutant was still lower than in the wild type at low density. This could explain the loss of the density effect showed in Artyukhin et al., 2013. Strikingly, there was a reactivation of DAF-16 in the *daf-6* mutant after 5 days in starvation (Fig. 3B, Supp. Fig. 2A). Although further investigations are necessary to explain this phenomena, it is possible that neuronal sensing is not completely abolished since some sensory cilia remain exposed in the mutant (Oikonomou et al., 2011). In conclusion, a correct functioning of the sensorial system was necessary for the density effect in arrested L1s, but the nature of the secreted signal remained still unknown.

### Trehalose present in the media during L1 arrest modulates DAF-16 activation

Arrested L1 larvae undergo a carbon metabolism shift towards the synthesis of the disaccharide trehalose promoted by DAF-16, which supports survival during starvation (Hibshman et al., 2017). Moreover, trehalose is able to activate DAF-16 in adult worms, since the addition of this sugar induces the expression of several DAF-16 targets (Seo et al., 2018). The evaluation of these two results together would suggest a positive feedback loop, where DAF-16 would promote the synthesis of trehalose and the disaccharide would, in turn, activate DAF-16. Therefore, we explored the involvement of trehalose as signaling molecule in the density effect on the L1 response to starvation. We first checked the possible presence of trehalose in the M9 medium where L1 larvae were arrested. To detect trehalose we used an enzymatic assay that involves the participation of trehalase, the enzyme that specifically hydrolyzes trehalose to two glucose molecules. We measured trehalose from the supernatant of L1 larvae arrested at different densities and we found trehalose in the µM range, which proportionally increased with the population density of the sample (Fig. 4A, left). We also found that trehalose concentration, normalized to the number of larvae hatched in each experiment, increased in the medium throughout the starvation period (Fig. 4A, right). There was a fast increase from 1 day after egg preparation until day 5, indicating that there is a secretion of trehalose by the larva to the medium during the first days of starvation. Then, trehalose concentration stabilized until the end of the experiment at day 15 of starvation. This suggests that there is not an effective consumption of the secreted disaccharide during the arrest, although we cannot discard an equilibrium between secretion and consumption after those first days of starvation. We then tried to elucidate the molecular response exerted by the addition of trehalose to larvae arrested at low density. For the following experiments we decided to use a trehalose concentration of 5 µM, which is the approximated concentration found in the supernatant of L1s arrested at high density, 20 L1/µl. The trehalose concentration measured for the DAF-16 reporter strain was slightly higher than for the *wt*, but not statically significant (Supp. Fig. 2B). Thus, exogenously added trehalose restored DAF-16 activation in worms at low density to similar levels to the larvae at high density (Fig. 4B, Supp. Fig. 3). This result was supported by the reduction in DAF-28 production when trehalose was added to L1s at low density (Fig. 4C). The insulin repression by trehalose was lost in the *daf-16* mutant, indicating the participation of DAF-16 in the blockage of ILP expression in sensory neurons in response to trehalose. As a consequence, the lack of *daf-16* also increased the synthesis of DAF-28 at high density when compared to wild type (Fig. 4C). These data reflect a positive feedback loop between DAF-16 and DAF-28, which was already known (Kaplan et al., 2019). We conclude that the trehalose secreted by the arrested L1s is sensed by the larvae, and that it influences the DAF-16/DAF-28 feedback, finally resulting in a higher DAF-16 activation. The diluted trehalose concentration in the medium of animals at low density results in a lower activation level of DAF-16.

**Figure 4.**
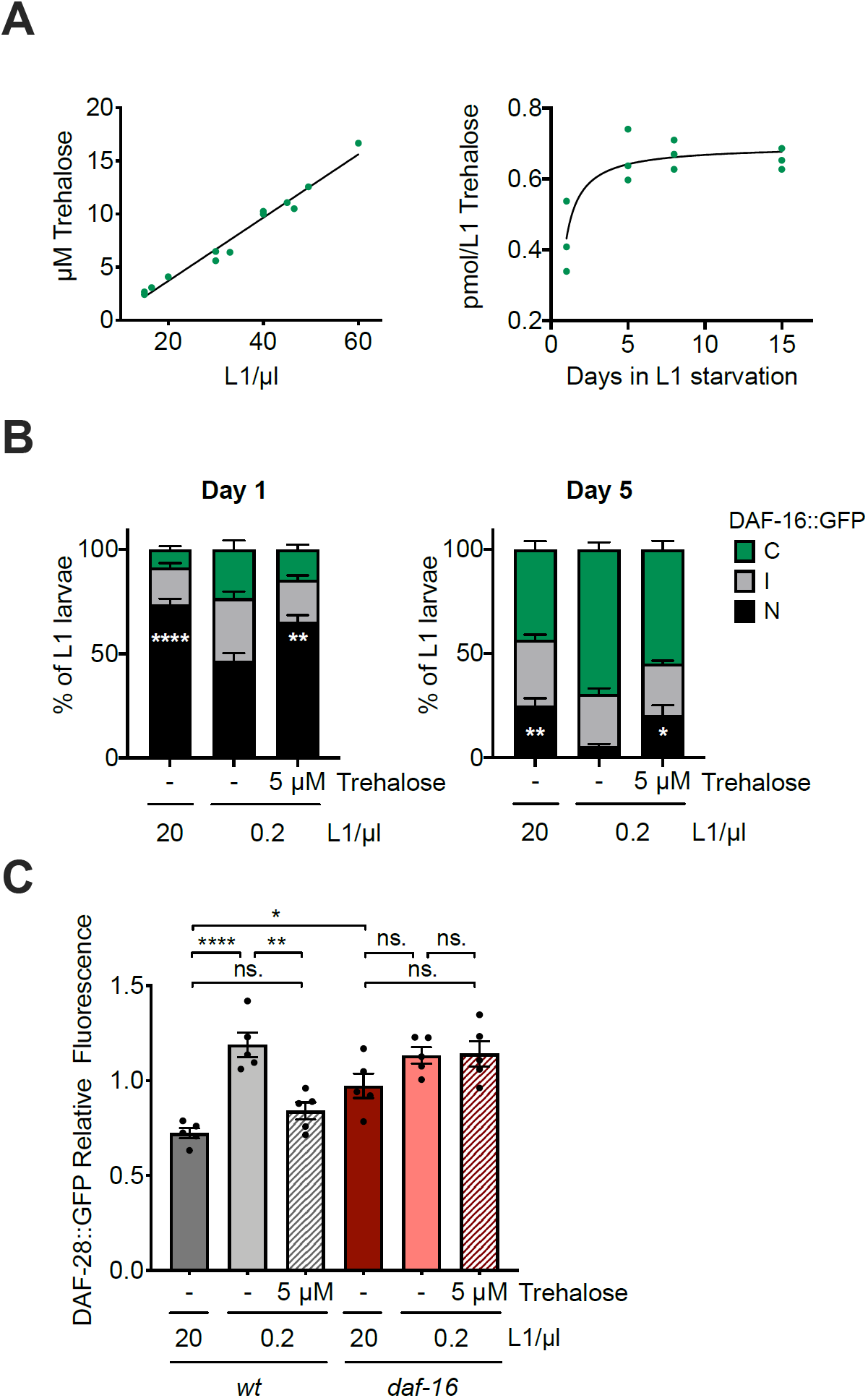
Secreted trehalose mediates DAF-16 activation in arrested L1 larvae. (A) Trehalose concentration (µM) in the medium of L1 larvae arrested at different densities after 1 day of starvation (left panel). Trehalose content in the medium during prolonged starvation, relative to the number of L1 larvae in the sample after one day of embryo preparation (right panel). (B) Subcellular localization of DAF-16::GFP after 1 and 5 days of starvation upon addition of 5 µM of trehalose to L1 larvae arrested at 0.2 L1/µl, compared with animals arrested at 20 and 0.2 L1/µl without trehalose. The histograms show the means of eight independent replicates and the error bars represent the SEM. Statistics for nuclear localization of DAF-16 are shown (* p<0.05, ** p<0.01, **** p<0.0001, one-way ANOVA). Independent plots for the three localization categories are depicted in Supp. Fig. 3. (C) DAF-28::GFP accumulation in the head of the animals in response to the addition of 5 µM of trehalose to L1 wild type and *daf-16* mutant larvae arrested at 0.2 L1/µl after 1 and 5 days of starvation, compared with animals arrested at 20 and 0.2 L1/µl without trehalose. Fluorescence measured for every animal was normalized to the mean signal of each independent replicate. Black dots represent the average relative fluorescence of each replicate. The histograms show the means of five independent replicates and the error bars represent the SEM. Averages of the replicates were used for statistics (* p<0.05, ** p<0.01, **** p<0.0001, one-way ANOVA).

### Trehalose does not reverse the detrimental effects of low density on its own

Since trehalose addition restored DAF-16 activation in arrested larvae at low density, we studied if it was also able to reverse the survival and recovery phenotypes associated to population density. When trehalose was added to the medium of larvae arrested at low density to a final concentration of 5 µM, the recovery percentage of those animals did not improve to high-density levels (Fig. 5A). Neither did the recovery time, although it showed a statistically non-significant shortening (Fig. 5B). These results indicate that trehalose, on its own, is not sufficient to mediate the density effect. However, we show here that density effect on the percentage of animals able to recover was completely dependent on DAF-16 and that trehalose addition was sufficient to reactivate DAF-16 in animals at low density. Two possible explanations would resolve this dilemma, both involving the participation of other signaling molecules. On one hand, trehalose could trigger a response in the treated animals, and potentially promote the secretion, in a DAF-16 dependent manner, of other molecules involved in the density effect. On the other hand, DAF-16 could be necessary but not sufficient to control the density effect, due to the necessity of other factors dependent on signals different from trehalose. In both cases, these putative signaling molecules would never reach a sufficient concentration in the medium of the worms at low density to reverse the phenotype when treated only with trehalose. However, in animals at high density, they could reach an optimal concentration, what would allow us to observe the actual participation of trehalose. To evaluate this possibility, we washed off the medium of L1 larvae arrested at high density after 24 hours of egg preparation, so every molecule released during hatching or secreted by the larvae were removed. Even then, larvae would continue secreting molecules to the medium, that eventually could reach a high concentration, so the expected phenotypes would not be as dramatic as in the animals at low density. Doing this, we first analyzed the recovery percentage of the worms after washing and how trehalose addition could alter it. We observed that recovery percentage of washed worms was somewhat reduced, and that this impairment was rescued by trehalose addition, however differences were not statistically significant (Fig. 6A). Since these were very long experiments, in which the animals could be actively secreting trehalose, we decided to use a mutant lacking the two trehalose synthases (TPS-1 and TPS-2) of *C. elegans*. This mutant was more sensitive to the wash, and the recovery percentage was rescued by the addition of trehalose (Fig. 6B). This result is in agreement with the possible presence of other secreted molecules that improve survival. The wash transiently reduced the level of DAF-16 activation, and it was recovered by trehalose addition, when we measured after 1 day of starvation (Fig. 6C, Supp. Fig. 4B). However, after 5 days of starvation this effect was no longer observable, and even DAF-16 activation was a bit higher in the washed sample. To further explore the role of trehalose as signaling molecule, we analyzed the DAF-16 activation level in the *tps-1; tps-2* double mutant. Here, we found that there were no differences in the DAF-16 localization between high and low densities, and its activation level was lower in the mutant than in the wild-type strain (Fig. 6D, Supp. Fig. 4C), what confirms that trehalose is necessary for the effect of density in DAF-16 activity.

**Figure 5.**
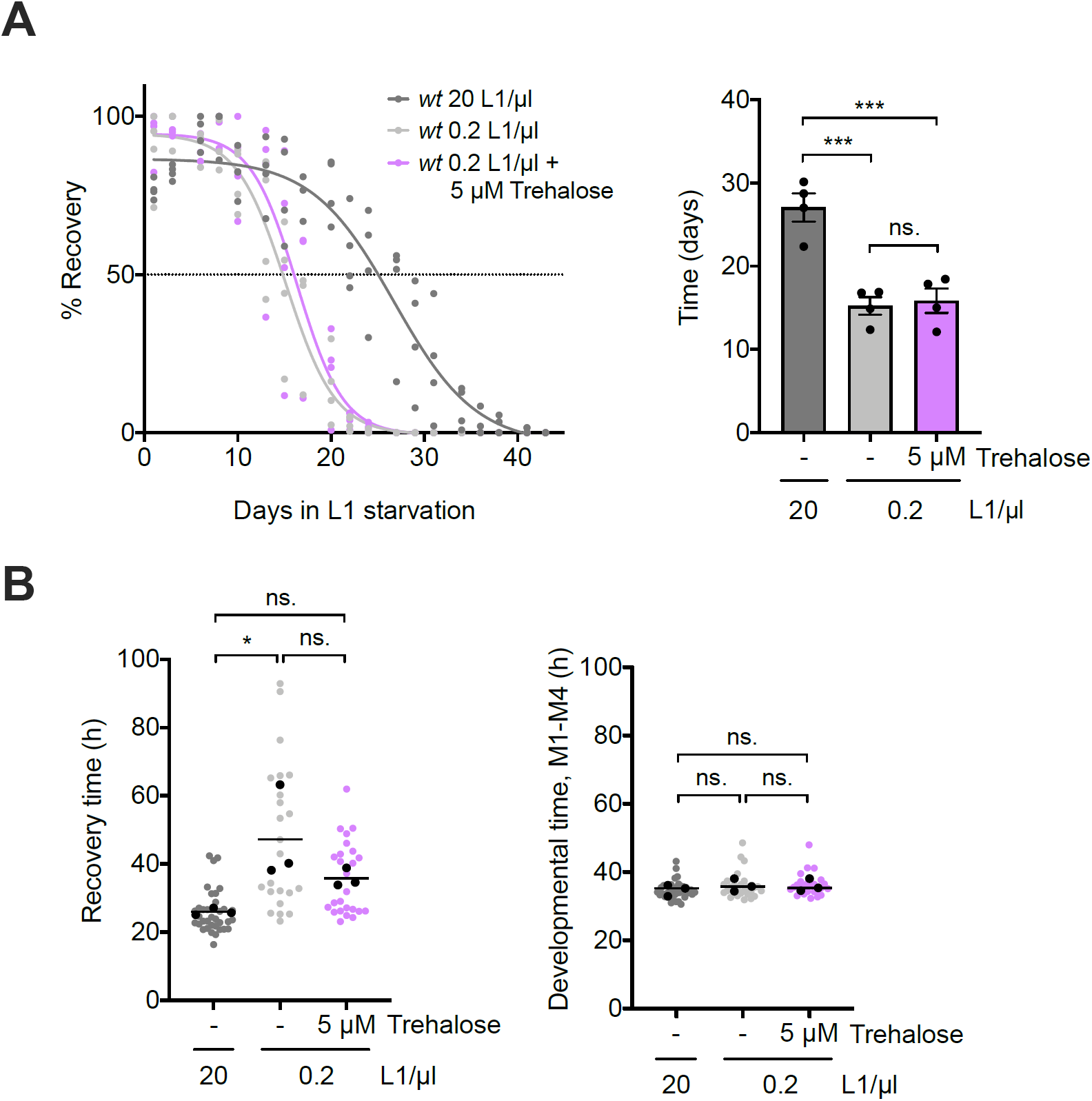
Trehalose supplementation is not sufficient to reverse survival and recovery of larvae arrested at low density. (A) Effect of the addition of 5 µM of trehalose on the percentage of wild-type animals arrested at 0.2 L1/µl that recover after different periods of starvation, compared with animals arrested at 20 and 0.2 L1/µl with no trehalose addition. Left: Percentage of recovery calculated every 2-3 days is independently plotted as colored dots for the four biological replicates. Curve fitting was performed using the data set from the four replicates. The horizontal dotted line marks the 50% recovery (half-life). Right: half-lives calculated from the fitting recovery curves. Black dots represent the value of each independent replicate. The bars mark the mean of the entire experiment and error bars shows the SEM (*** p<0.001, one-way ANOVA). (B) Recovery time (L1) and developmental time (M1-M4) after 8 days of starvation of animals arrested at 0.2 L1/µl treated with trehalose 5 µM and animals arrested at 20 and 0.2 L1/µl without trehalose. Every colored dot represents the data from each animal analyzed. Black dots represent the average of each of the three independent replicates and the line marks the mean of the entire experiment. Averages of the replicates were used for statistics (* p<0.05, one-way ANOVA).

**Figure 6.**
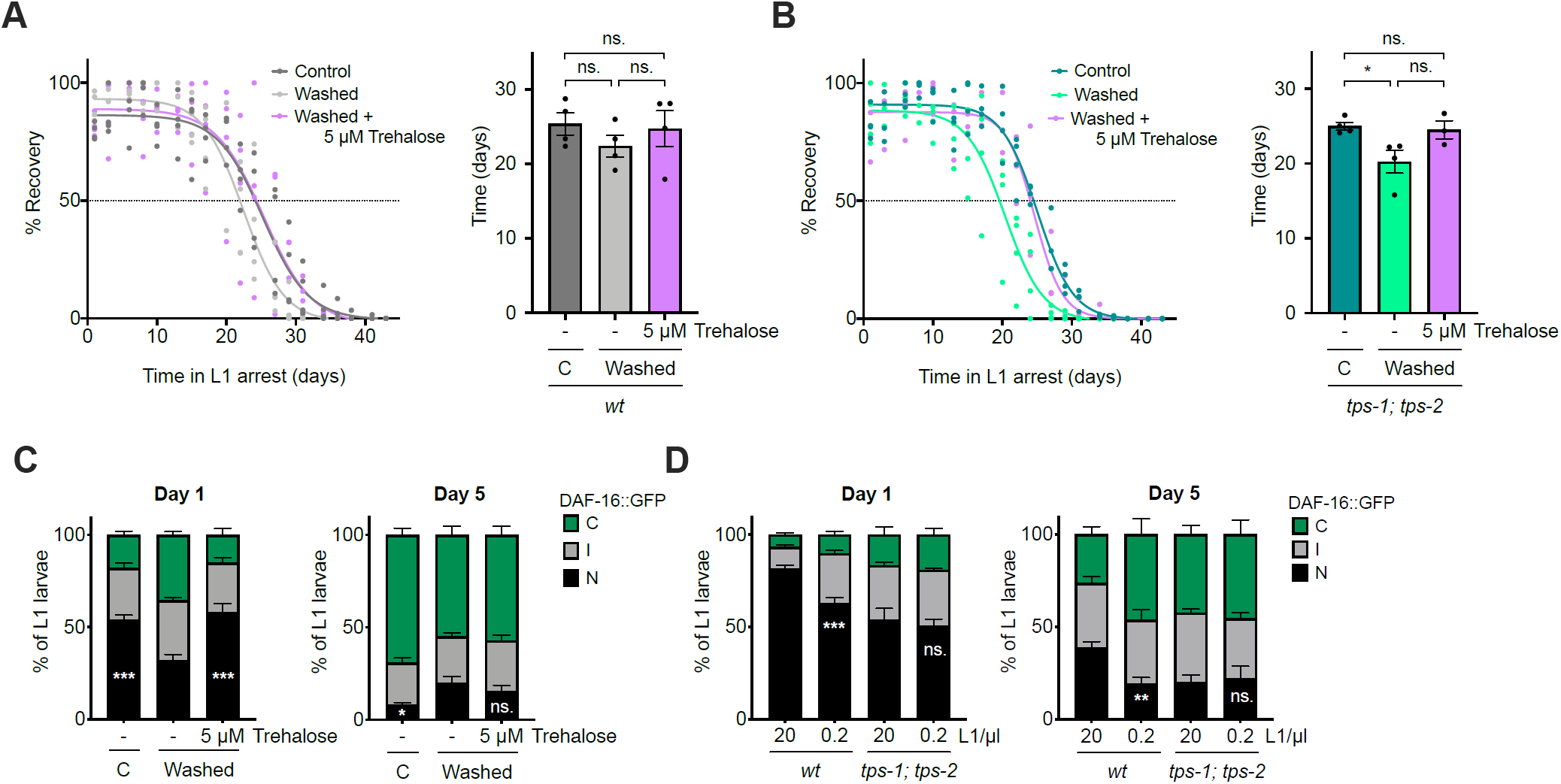
Trehalose mediates the response in animals at high density. Effect of trehalose addition (5 µM) on the percentage of wild-type (A) and *tps-1; tps-2* (B) animals that recover from different periods of arrest at 20 L1/µl after washing off the medium where larvae hatched. In control samples, medium was not washed and trehalose was not added. Recovery curves for wild-type (A) and *tps-1; tps-2* mutant (B) are plotted in the corresponding left panels. Each percentage calculated every 2-3 days is independently plotted as colored dots for the four biological replicates. Curves fitting was performed using the data set from, at least, three replicates. The horizontal dotted line marks the 50% recovery (half-life). In the right panels, half-lives calculated from the fitting curves in the left panels. Black dots represent the average of each independent replicate. The bars mark the mean of the entire experiment and error bars shows the SEM. Averages of the replicates were used for statistics (*** p<0.001, one-way ANOVA). (C) Subcellular localization of DAF-16::GFP in L1 larvae arrested at 20 L1/µl at 1 and 5 days after washing the medium. Trehalose was added to one sample to a final concentration of 5 µM. In control sample the original medium was not removed and trehalose was not added. (D) Subcellular localization of DAF-16::GFP in wild type and *tps-1; tps-2* L1 larvae arrested at 20 and 0.2 L1/µl after 1 and 5 days of starvation. (C and D) The histograms show the means of, at least, four independent replicates and the error bars represent the SEM. Statistics for nuclear localization of DAF-16 are shown (* p<0.05, ** p<0.01, *** p<0.001, one-way ANOVA in (C) and Unpaired t-test in (D)). Independent plots for the three localization categories are depicted in Supp. Fig. 4B and C.

This process must involve a receptor that senses exogenous trehalose. We tested GUR-3 as possible trehalose receptor in *C. elegans*, since this is the only protein in the worm with homology to Gr5a, the trehalose receptor of *Drosophila melanogaster* (Chyb et al., 2003). GUR-3 is a gustatory receptor expressed in different neurons, including the pharyngeal neurons I2s, and it is involved in the sensing of hydrogen peroxide, light and temperature (Bhatla and Horvitz, 2015; Goya et al., 2016). GUR-3 and Gr5a have a global similarity of 30% and an identity of 14.9%. However, according to the domain prediction tool SMART, GUR-3 conserves the trehalose receptor domain of Gr5a (Supp. Fig. 5 A and B). To check if the *gur-3* mutant was sensitive to the presence of trehalose in the medium, we analyzed the DAF-16 activation level at low density and high density, and in response to the addition of trehalose. Unlike what happens in the *wt*, neither density nor trehalose affected the nuclear localization of DAF-16 in the *gur-3* mutant. Overall, nuclear DAF-16 in *gur-3* mutants was reduced compared to wild type animals (Fig. 7, Supp. Fig. 5 C). This indicated that the *gur-3* mutant was insensitive to the presence of trehalose, supporting a role for GUR-3 as a trehalose receptor.

**Figure 7.**
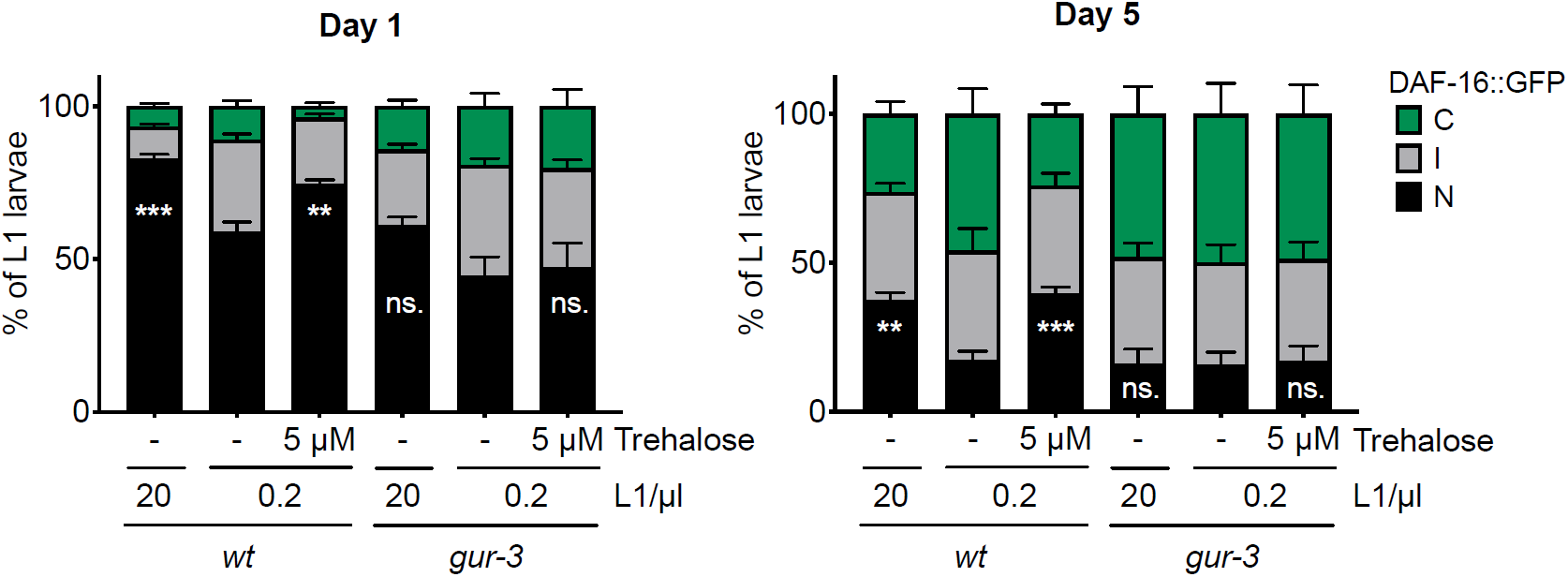
GUR-3 mediates DAF-16 activation by trehalose addition. Subcellular localization of DAF-16::GFP in wild type and *gur-3* L1 larvae arrested at 20 and 0.2 L1/µl after 1 and 5 days of starvation. Trehalose to a final concentration of 5 µM was added to animals at low density. Animals at high density were used as control. The histograms show the means of, at least, three independent replicates and the error bars represent the SEM. Statistics for nuclear localization of DAF-16 are shown (** p<0.01, *** p<0.001, one-way ANOVA). Independent plots for the three localization categories are depicted in Supp. Fig. 5 C.

## DISCUSSION

Social interaction plays a protective role in arrested L1s. High density population improves survival of arrested L1 larvae to starvation (Artyukhin et al., 2013a). Aggregation behavior displayed by arrested L1s on agar plates in absence of food generates a local high density of animals, which could promote starvation survival (Artyukhin et al., 2015). Aggregation is also beneficial against external stressors in adult *C. elegans* (Boender et al., 2011). Longevity is extended when exposed to low oxygen environments, below normoxia (21%) (Honda et al., 1993). Aggregation provides to *C. elegans* its preferred oxygen levels (6 %) inside the aggregate (Gray et al., 2004; Rogers et al., 2006). This behavior also occurs in response to other noxious stimuli (Boender et al., 2011). Moreover, the high-density population in the aggregate leads to *dauer* formation, which allows worms to survive until the environmental conditions are optimal (de Bono et al., 2002). Therefore, it appears that *C. elegans* has adopted a collective social behavior entailing high population density as strategy to resist detrimental conditions.

Here we show that high population density not only enhances survival of arrested L1 to starvation (Artyukhin et al., 2013a), but it also predisposes to a better recovery after arrest (Fig. 3A). This means that these animals will develop and adapt faster to new optimal conditions. Thus, in a situation of competition for the same resources with other species, *C. elegans*, as population, will be able to optimize the available resources and will have an advantage. In the case of *Caenorhabditis* species with no social behavior, individual animals adopt an explorative behavior in absence of food, and, as a consequence, they lack the starvation survival enhancement in response to high-population density (Artyukhin et al., 2015).

With the aim of unravelling the molecular mechanisms underneath, we have tested DAF-16, a main determinant of L1 starvation survival (Baugh and Sternberg, 2006; Lee and Ashrafi, 2008; Muñoz and Riddle, 2003) and recovery (Olmedo et al., 2020). Arrested L1s are more resistant to stress (Muñoz and Riddle, 2003; Weinkove et al., 2006), since the activation of DAF-16 during starvation leads to up-regulation of stress response genes (Baugh et al., 2009; Kaplan et al., 2019). DAF-16 activation directly influences starvation survival, but also determines the level of accumulation of signs of aging during prolonged L1 arrest. Thus, arrested *daf-2* mutant L1 larvae, which present higher DAF-16 activation, display lower levels of aging phenotypes after long periods of starvation (Olmedo et al., 2020). The progressive accumulation of these phenotypes during the aging process in arrested L1s impairs their recovery capacity after refeeding (Olmedo et al., 2020; Roux et al., 2016). Here we show that the density effect on the capacity of arrested L1s to recover from prolonged starvation depends on DAF-16, whose activation is higher at high densities (Fig. 2E). This might be interpreted as if the lower DAF-16 activation level at low density, and its absence in the mutant, leads to a lower stress resistance response. This reduced response could lead to a higher accumulation of cellular damage during prolonged arrest, resulting in an impaired capacity for recovery, and reduced survival. We would like to note that the effect of lacking DAF-16 is more severe than the effect of being arrested at low density in terms of percentage of recovery (Fig. 2A and B). This is also evidenced by the fact that, while none of the *daf-16* L1 larvae recovered after 8 days of arrest, a number of wild-type animals arrested at low density did recover, allowing us to estimate recovery time (Fig. 1B). This suggests some DAF-16 protective activity in worms at low density, which is completely absent in the mutant. This can also be seen in the level of DAF-16 activation, which is detected even at low density (Fig. 2D, Fig. 3B, Fig. 4A, Fig. 6D and Fig. 7). Total absence of DAF-16 activation is only observed in the case of the *tax-4* mutant strain, which remains insensitive to density changes due to a sensory defect resulting in a constitutive activation of the IIS. Therefore, the density effect on recovery from starvation and survival requires DAF-16, which becomes more activated in animals that sense high-density cues.

The presence of unknown signaling molecules in the supernatant of arrested larvae supports survival in L1s at high density and recovers the short longevity of animals at low densities (Artyukhin et al., 2013a). According to our results, the conditioned medium from L1s arrested at high density was also able to rescue the longer recovery time of L1s at low density (Fig. 3A). So, these molecules are sufficient to trigger the resistance response to starvation, which allows the arrested L1s to survive longer and recover faster. These signaling molecules, abundant at high density, reduced the production of the insulin DAF-28 at the beginning of the starvation period (Fig. 2E). This reduction of insulin in sensory neurons would lead to an activation of DAF-16 (Kaplan et al., 2019). At the same time, DAF-16 activation seems to feed back to repress DAF-28 (Fig. 4C), since the *daf-16* mutant shows increased signal of DAF-28::GFP. This is consistent with previously published data. *C. elegans* contains 40 genes coding for insulin-like peptides (ILPs), which can act as agonist or antagonist of the insulin receptor DAF-2 (Kaplan et al., 2019; Pierce et al., 2001). Positive and negative feedback via DAF-16 regulation has been reported for most of the insulin peptides over L1 arrest, and in the transition to development resumption (Kaplan et al., 2019). DAF-28 is an agonist ILP, and as such it promotes development and reproductive growth in response to feeding (Chen and Baugh, 2014; Li et al., 2003). DAF-28 is repressed by DAF-16 in the arrested L1, suggesting a positive feedback (Kaplan et al., 2019). *daf-28* expression is downregulated during starvation or upon application of *dauer* pheromone (Li et al., 2003). Furthermore, it is required for the response to food perception, even when ingestion is absent (Kaplan et al., 2018), and its expression also decreases in response to high temperature (Kaplan et al., 2019). Therefore, *daf-28* expression is regulated by a variety of environmental cues, including molecules in the medium and temperature. So, it appears reasonable that the production of this insulin in the sensory neurons could be regulated by the secreted molecules that mediate communication between animals.

In a search for possible communication molecules that could be involved in the effect of density, we tested trehalose. We chose it because it is a molecule whose synthesis is upregulated in arrested L1s (Hibshman et al., 2017) and it activates DAF-16 (Seo et al., 2018). Trehalose is present in a wide variety of organisms (Elbein et al., 2003). It is capable of stabilizing membranes and proteins, what makes it a protective molecule against a diversity of stresses, including desiccation, heat, cold, and oxidative stress. Trehalose also prevents denatured proteins from aggregating (Crowe, 2007; Elbein et al., 2003; Singer and Lindquist, 1998). In *C. elegans*, trehalose is necessary for desiccation tolerance of the *dauer* larvae (Erkut et al., 2011). Trehalose addition to the adults enhances longevity and thermotolerance (Honda et al., 2010). This longevity extension is completely dependent on DAF-16 and autophagy, both activated by trehalose (Seo et al., 2018). In arrested L1 larvae, trehalose supports starvation survival (Hibshman et al., 2017). However, here we tested for a different role of trehalose, as a possible signaling molecule that would determine the effect of density. A similar role has been described in plants, where trehalose-6-phosphate works as signal of sugar availability and regulates responses to starvation (Elbein et al., 2003; Smeekens et al., 2010). To work as a communication signal, trehalose should be free in the medium, transiently at least. In fact, we detected the presence of trehalose in the media of arrested wild-type and the DAF-16::GFP reporter strain (Supp. Fig. 2B). Importantly, the concentration of trehalose was proportional to the number of larvae hatched and it was in the µM range (Fig. 4A). Internal trehalose concentration previously determined in L1 larvae was in the range of 100 nmol/mg protein in both starved and fed animals (Hibshman et al., 2017). This amount was already present in the embryos before hatching and increased within the 12 hours after hatching in the starved animals, meanwhile it decreased in the fed ones (Hibshman et al., 2017). Our measurements are not comparable, since we analyzed the trehalose in the supernatant and they measured the trehalose present in the whole animal. A concentration higher than 20 mM is needed to see the protective effect of trehalose during the L1 starvation, being 50 mM the concentration used for most of those experiments (Hibshman et al., 2017). That represents a concentration 10,000 times higher than that detected in the supernatant. Therefore, it appears that, for trehalose to play its direct role as stress protectant, it must be in large quantities. Additionally to its protective role, trehalose works as energy source when added at high concentration, since it has to be hydrolyzed to glucose by the action of the trehalase enzymes in order to observe its complete beneficial effect on survival (Hibshman et al., 2017). However, what we propose here is a signaling role for trehalose, where it works at low concentration, being sensed by the animals and then triggering a response. Our data suggests that, for that role, trehalose does not need to be consumed, as we observe an increase in the amount of trehalose present in the supernatant of arrested L1s over the time in starvation (Fig. 4A). Furthermore, the small amount of trehalose that we measured in the supernatant was enough to reduce DAF-28 production and to activate DAF-16 (Fig. 4B and 4C). This response is contrary to what is expected for a sugar that works as carbon and energy source, as it happens in the case of glucose (Lee et al., 2009) or when food is perceived (Kaplan et al., 2018). In these cases, insulin signaling is enhanced and DAF-16 inactivated. Therefore, the role of trehalose described here is different to other roles as protectant and as energy source.

No trehalose content was previously detected in L1 larvae of the double *tps-1; tps-2* mutant, which lacks both trehalose synthases (Hibshman et al., 2017). Here, in the supernatant of the double *tps* mutant we detected about a third of the trehalose detected for the wild type (Supp. Fig. 4A). The trehalose reduction detected in the double mutant was sufficient to get reflected in a higher DAF-16 inactivation compared to the wild type (Fig. 6D, Supp. Fig. 4C). We cannot be sure about the origin of this trehalose in the mutant, but we hypothesize that it was ingested by the mother with the food, then provided to the embryos and finally released during hatching. Trehalose is present at high concentration in the embryos of many different nematode species, and its participation is determinant for embryo development and for the hatching process (Behm, 1997). During hatching, the permeability of the innermost layer of the eggshell increases, which results in the leakage of trehalose and other metabolites (Behm, 1997; Perry, 1989). Two protective roles of trehalose have been proposed in nematode eggs. One role would involve the stabilization of the lipid membrane of the eggshell, which would allow the resistance of the embryos to extreme environments. The second one would involve the release of trehalose to the surrounding medium, what would reduce the osmotic stress (Perry, 1989). However, although trehalose was found in *C. elegans* embryos before hatching (Hibshman et al., 2017), the function of trehalose in embryos of this nematode remains to be demonstrated. Nevertheless, it has been shown that *C. elegans* mothers can adapt the sugar provisioning to the embryos in response to changing environmental conditions (Dey et al., 2016; Frazier and Roth, 2009). The content in trehalose, glycogen and glycerol changes in the embryos of mothers exposed to different stresses. More specifically, trehalose provisioning increased in embryos from mothers exposed to hyperosmotic stress (Frazier and Roth, 2009). Despite the relevance of trehalose in embryos, and the importance of maternal provisioning, it remains unexplored whether trehalose ingested by the mother (as opposed to synthesized by the worm) can be used to provide the embryos. A trehalose transporter in *C. elegans* has not been identified yet. However, in mammals, a glucose carrier is able to transport trehalose (Mayer et al., 2016). This could be a possibility to investigate in *C. elegans*, being the glucose carrier FGT-1 the closest homolog to the trehalose transporter Tret1-1 in *Drosophila*. Actually, the reduction in the expression of *fgt-1* leads to an extended longevity, not additive to the extension displayed in *daf-2* and *age-1* mutants, meaning that insulin signaling is reduced when *fgt-1* expression is down (Feng et al., 2013).

To work as a signaling molecule, trehalose must be sensed by the animals. Here we tested GUR-3, the closest homolog to Gr5a in *C. elegans*, as a possible trehalose receptor. Gr5a is a G protein-coupled gustatory receptor of *Drosophila* required to activate behavioral and sensory responses to trehalose (Chyb et al., 2003). Trehalose sensing in *Drosophila* regulates lifespan and the mutant lacking Gr5a lives shorter (Waterson et al., 2015). In flies, trehalose is the most abundant circulating sugar, and in the *gr5a* mutant there is an increased trehalose synthesis in an attempt to compensate for the lack of trehalose sensing (Waterson et al., 2015). This is a clear example of trehalose functioning as a signal molecule rather than a protective molecule. In our experiments we observed that the *gur-3* mutant is insensitive to trehalose in terms of DAF-16 activation (Fig. 7). However, further investigation is needed to corroborate the capacity of GUR-3 to sense trehalose in the medium and trigger the subsequent response. For instance, one of the questions to be addressed concerns the localization of GUR-3 in neurons different to ASI and ASJ, the sensory neurons where DAF-28 is mainly expressed in the head of the worms (Li et al., 2003). This seems to be important since the trehalose signaling that we propose here involves inhibition of the DAF-28 production (Fig. 4C) and probably of other ILPs. Thus, new studies could reveal the connection between these neurons or the location of the GUR-3 receptor in more neuronal types to the ones already described. However, different mechanisms or receptors cannot be completely discarded at this point. An example of alternative mechanism is what was found in mammalians where trehalose binds and blocks the glucose carriers SLC2A to induce autophagy (DeBosch et al., 2016).

Although low concentrations of trehalose activated DAF-16 in our experiments, this response was not sufficient to support starvation survival and recovery of animals arrested at low density (Fig. 5A and B). This makes us think that trehalose could work triggering the response only at the first moments of starvation and that the participation of other molecules would be then necessary. Actually, even when trehalose secretion continues over the arrest period (Fig. 4A), we only observed the effect of density on DAF-28 production after 1 day of arrest, but not on day 5 (Fig. 2E). Moreover, DAF-16 also becomes less active throughout prolonged starvation, independently of the density (Fig. 2D). The DAF-16 inactivation during long-term starvation had been previously described and it requires the activity of AGE-1, in a DAF-2 independent manner (Weinkove et al., 2006). This would decouple the inactivation of DAF-16 from the presence of trehalose after the first days of the prolonged starvation. Therefore, the role of trehalose in this response would be restricted to the first days of arrest, probably due to the later activation of additional mechanisms or pathways, such as AGE-1, which would limit the response to trehalose. Consistent with the idea that additional molecules are involved in the density effect, we found recovery percentage enhancement by trehalose addition in larvae starved at high density where the original medium containing the secreted molecules was removed. This was clear in the double *tps* mutant (Fig. 6B), since there was not active trehalose production, despite some trehalose possibly contained in the embryos, as previously discussed.

The synthesis of a disaccharide in the absence of food appears to be paradoxical, but trehalose is known to be protective, and as such it acts on arrested L1 larvae, where it supports survival (Hibshman et al., 2017). It also seems clear that it can be used as an energy source as a last resort. However, what we propose here is an additional role in the communication between animals. On this role, trehalose would trigger the response against stress in the animal that senses the signal. This response involves the IIS pathway, which gets inactivated in the first moments of prolonged starvation, supporting starvation survival. However, to maintain the effect on survival and recovery, the participation of other unknown secreted molecules is needed. Nevertheless, this resistance mechanism is lost in animals arrested at low density, since secreted signals are diluted, and the response is not activated sufficiently. Therefore, in order to resist environmental stresses, remaining as high-density population is beneficial for *C. elegans*.

## MATERIAL AND METHODS

### *C. elegans* strains and growth conditions

We cultured and maintained animals according to standard methods (Brenner, 1974), typically at 20 °C on nematode growth medium (NGM) plates seeded with a lawn of *Escherichia coli* OP50-1, used as nematode nutrient source. The N2 Bristol strain background is referred in the text as wild type. We performed longevity assays on N2, CF1038 *daf-16(mu86) I*, and MOL273 *tps-1(ok373) X; tps-2(ok526) II*. We generated MOL273 using routine genetic techniques by crossing VC225 *tps-1(ok373) X* and RB760 *tps-2(ok526) II*. To quantify recovery time by luminometry we used the reporter strain MRS387 *sevIs1 [Psur-5::luc+::gfp] X* (Olmedo et al., 2020). To determine the subcellular localization of DAF-16 we used the strain TJ356 *zIs356 [Pdaf-16::daf-16a/b-gfp; rol-6] IV*. We crossed TJ356 with CB1377 *daf-6(e1377) X*, PR678 *tax-4(p678) III*, MOL273 *tps-1(ok373) X; tps-2(ok526) II* and RB1755 *gur-3(ok2245) X* to generate MOL274 *daf-6(e1377) X; zIs356[Pdaf-16::daf-16a/b-gfp; rol-6] IV*, MOL216 *tax-4(p678) III; zIs356[Pdaf-16::daf-16a/b-gfp; rol-6] IV*, MOL271 *tps-1(ok373) X; tps-2(ok526) II; zIs356[Pdaf-16::daf-16a/b-gfp; rol-6] IV* and MOL276 *gur-3(ok2245) X; zIs356[Pdaf-16::daf-16a/b-gfp; rol-6] IV*. We used the strain VB1605 *svIs69[Pdaf-28::DAF-28::GFP]* to quantify DAF-28 expression in the head of the arrested larvae. We crossed this strain with CF1038 to generate MOL224 *daf-16(mu86) I; svIs69[Pdaf-28::DAF-28::GFP]*. The strains not generated here were obtained from the Caenorhabditis Genetics Center (CGC).

To arrest L1 larvae by starvation we first obtained embryos from gravid adults by treatment with alkaline hypochlorite solution. Those embryos were incubated in M9 minimum medium at 20°C with gentle shaking leading to their hatching. These larvae newly hatched in absence of food arrest their development as L1. The density of the worm culture was adjusted to the indicated in every experiment after embryo preparation. Although after hatching the population density may vary from the calculated, we considered it important to adjust immediately after embryo preparation, so that the larvae already hatched at the approximate desired density and the released compounds they encountered are consequently at high or low concentration. For recovery experiments depicted in Fig. 1, where different starvation times were applied, we staggered the embryo preparation to be able to simultaneously analyze the recovery of animals with different periods of arrest.

For the experiments in which we washed off the medium where the animals hatched, we spun down twice the samples of arrested L1s during 2 min at 850 g and resuspended the animals into clean M9. In parallel, we also centrifuged the controls, but, in these cases, animals were not resuspended in clean M9 but in their original medium.

To prepare the conditioned medium (CM), we obtained embryos and adjusted the sample density at higher than 50 embryos/µl. The day after we counted the number of hatched animals so we could adjust the arrested L1s to a final concentration of 50 L1/µl. We then centrifuged them twice during 10 min at 5000 g to collect the supernatant without larvae. We froze this CM at −20 °C until the moment of use.

### Determination of recovery time

Time needed to recover after prolonged periods of arrest was measured using a bioluminescence-based method that allows the precise quantification of every larval and molt stage during the post-embryonic development (Olmedo et al., 2015). Briefly, arrested L1 larvae were individually pipetted into a well of a white 96-well plate containing 200 µM Luciferin in 100 µl of S-basal (including 10 µg/ml of cholesterol prepared in ethanol). Once all animals were placed into the plate, we added the food to resume development. Thus, we pipetted into every well 100 µl of S-basal containing 20 g/l of *E. coli* OP50-1. Prior to measuring, we sealed the plates with a gas-permeable cover (Breath Easier, Diversified Biotech). Then, luminescence of each animal was recorded by a highly sensitive microplate luminometer (Berthold Centro LB960 XS^3^) for 1 s at intervals of 5 min until animals reached adulthood. Luminometers were kept inside temperature-controlled incubators (Panasonic MIR-154) to maintain a constant 20 °C temperature during the experiments. We used sample sizes of more than 20 animas per condition and experiment in, at least, three biological replicates. The luminometry data for each animal was analyzed individually and, for statistics, we used the averages of independent replicates to avoid the inflated N value from using individual animals. The analysis was performed as previously described (Olmedo et al., 2015). We use a template in MS-Excel for analysis of the luminescence signal and GraphPad Prism for statistics and graphs. The data collected by this technique allowed the calculation of the recovery time (L1) (Fig. 1B), the developmental timing (M1 to M4) (Supp. Fig. 1A) and the complete the post-embryonic development (L1 to M4) (Supp. Fig. 1B) for every animal analyzed.

### Analysis of the recovery percentage

To score the survival capacity to starvation of the arrested L1 larvae we calculated the percentage of animals able to recover when fed after prolonged starvation. To do that, we pipetted every 2-3 days a fix number of L1s arrested at the indicated densities onto NGM plates with food and after 2 days we counted the number of animals that were alive and did progress in development. We monitored the plates for few days more in case that some animals needed more time to recover, especially after long periods of starvation. The percentage of recovery on each day was calculated relative to the day with the maximum number of worms recovered for each strain or condition. This maximum was taken as the 100 %. We performed, at least, three independent biological replicates per experiment. All replicates were plotted together, and the medium survival curves were fit to these data. The half-life of each biological replicate was independently calculated for statistics. Curve fitting and half-lives were calculated using the analysis tools available in GraphPad Prism software.

### Subcellular localization of DAF-16

To determine the subcellular localization of DAF-16 we used the strains expressing the fluorescently tagged version of DAF-16 (DAF-16::GFP) described above. At the indicated times, we took aliquots of 18 µl of the suspensions of arrested L1 larvae and we placed them on a microscope slide with 2 µl of 10 mM Levamisole to paralyze the animals. We used a Leica M205 FCA stereo microscope to visualize DAF-16::GFP. Thus, we counted the number of animals with nuclear, cytoplasmatic or intermediate localization of DAF-16 in a population of, at least, 100 animals per sample. With these data, we calculated the percentage animals with each localization of DAF-16. We performed a minimum of 3 independent biological replicates. To avoid the artificial translocation of DAF-16 we used a reduced concentration of Levamisole. In the same way, during the visualization the room temperature was kept at 20 °C to avoid the possible effect of temperature in DAF-16 activation.

### DAF-28::GFP quantification

We used strains containing a translational GFP reporter of DAF-28 to quantify its expression in the head of the arrested L1 larvae. We took aliquots of 10 µl from suspensions of L1s arrested to the indicated densities after 1 or 5 days of arrest and we placed them on a microscope slide with 1 µl of 10 mM Levamisole to paralyze the animals. We then acquired pictures of these worms using a Leica M205 FCA stereo microscope. We analyzed those pictures with FIJI software, which allowed us to measure the signal intensity of a selected area. We analyzed more than 20 animals per condition and experiment, and we performed a minimum of 3 individual biological replicates per experiment. Due to discrepancies in the raw data between replicates, we normalized the signal measured for each worm to the average signal of the replicate. We used the averages of the normalized signal per condition and replicate to perform the statistics analysis.

### Trehalose measurement

Trehalose content in the medium of arrested L1 larvae was measured by enzymatic determination of released glucose after hydrolysis of trehalose by trehalase enzyme, using the glucose oxidase/peroxidase method (Sigma catalog #GAGO-20). We spun down the L1 cultures for 5 min at 5000 g to remove the animals and we collected the supernatant for measurement. We added potassium phosphate buffer pH 6 to the samples to a final concentration of 0.1 M and trehalase enzyme (SIGMA) to 1.5 U/ml. This mix was incubated overnight at 37 °C with no shaking. We then added the glucose oxidase/peroxidase mix containing 10 U/ml glucose oxidase, 2 U/ml horseradish peroxidase and 80 µg/ml O-dianisidine. We incubated the samples at 37 °C for 30 min, we then added H_2_SO_4_ at a final concentration of 4.3 N and we measured the OD of the samples at 540 nm. A standard curve was prepared in parallel with known concentrations of trehalose, so we could transform OD values into trehalose content. For each sample we performed a parallel reaction in which trehalase was substituted by water in order to detect free glucose, what would make us overestimate the content in trehalose. We referred the trehalose content measured to the number of larvae actually hatched for each sample, what we determined the day after egg preparation.

## ACKNOWLEDGMENTS

Some strains were provided by the *Caenorhabditis* Genetics Center (CGC), which is funded by NIH Office of Research Infrastructure Programs (P40 OD010440). We thank Ryan Baugh (Duke University) for helpful discussion. M.O is supported by the Ramón y Cajal program of the Spanish Ministerio de Economía y Competitividad, (RYC-2014-15551). Our work is supported by the Spanish Ministerio de Economía y Competitividad (BFU2016-74949-P), the European Research Council (ERC-2011-StG-281691) and a Marie-Curie Intra-European Fellowship (FP7-PEOPLE-2013-IEF/GA Nr: 627263).

## Supplementary material for

**Supplementary Figure 1.**
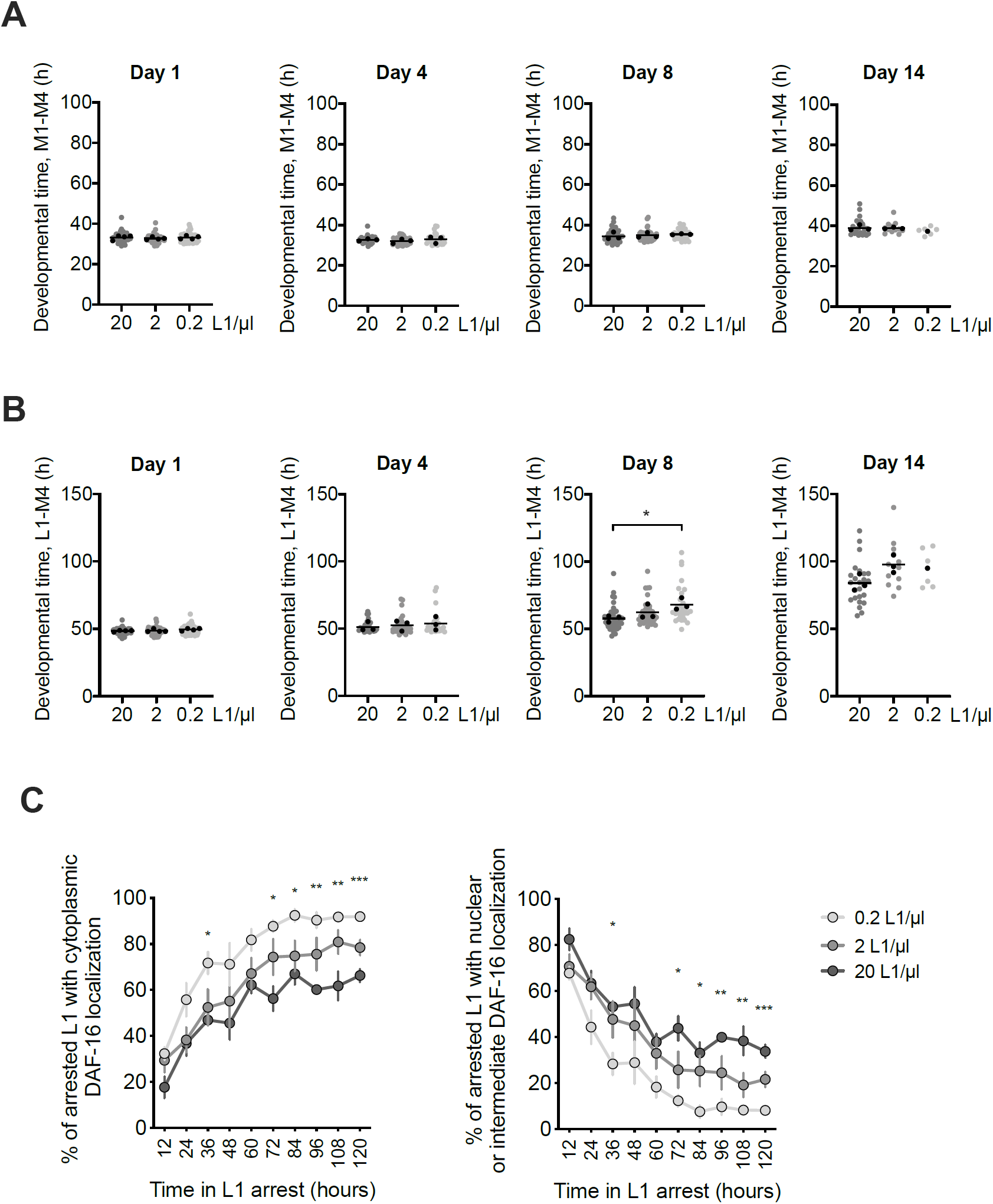
Developmental time (M1 to M4) (A) and complete the post-embryonic development (L1 to M4) (B) of animals arrested at densities of 20, 2 and 0.2 L1/µl, after 1, 4, 8 and 14 days of starvation. All individual animals from three independent biological replicates are plotted. Black dots represent the average of each independent replicate and the black line indicates the mean of the entire experiment. Averages of the replicates were used for statistics (* p<0.05, Unpaired t-test). (C) Subcellular localization of DAF-16::GFP in L1 larvae arrested at densities of 20, 2 and 0.2 L1/µl for 1 to 5 days in starvation. Dots indicate the mean of four independent replicates and the error bars represent the SEM (* p<0.05, ** p<0.01, *** p<0.001, one-way ANOVA).

**Supplementary Figure 2.**
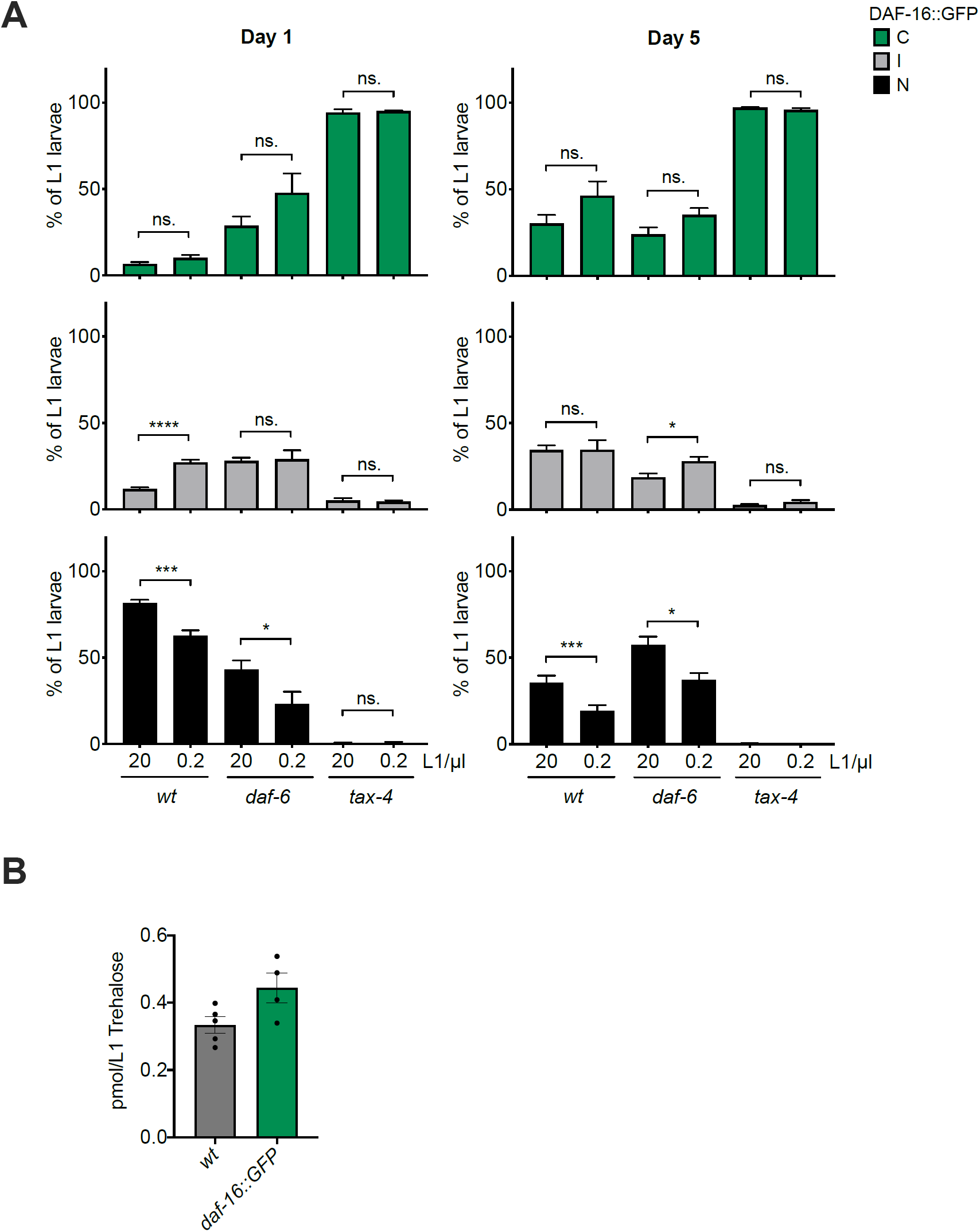
(A) Subcellular localization of DAF-16::GFP in L1 larvae arrested at 20 and 0.2 L1/µl for 1 and 5 days in *wt, daf-6* and *tax-4* mutant strains. Percentage of animals with cytoplasmatic localization are shown in the top panel, percentage of animals with intermediate localization are in the middle panel and nuclear localization of DAF-16 is shown in the bottom panel. The histograms show the means of, at least, three independent replicates and the error bars represent the SEM (* p<0.05, *** p<0.001, **** p<0.0001, Unpaired t-test). (B) Relative trehalose content per hatched animal in the medium of wild-type and TJ356 reporter strain, which expresses DAF-16::GFP.

**Supplementary Figure 3.**
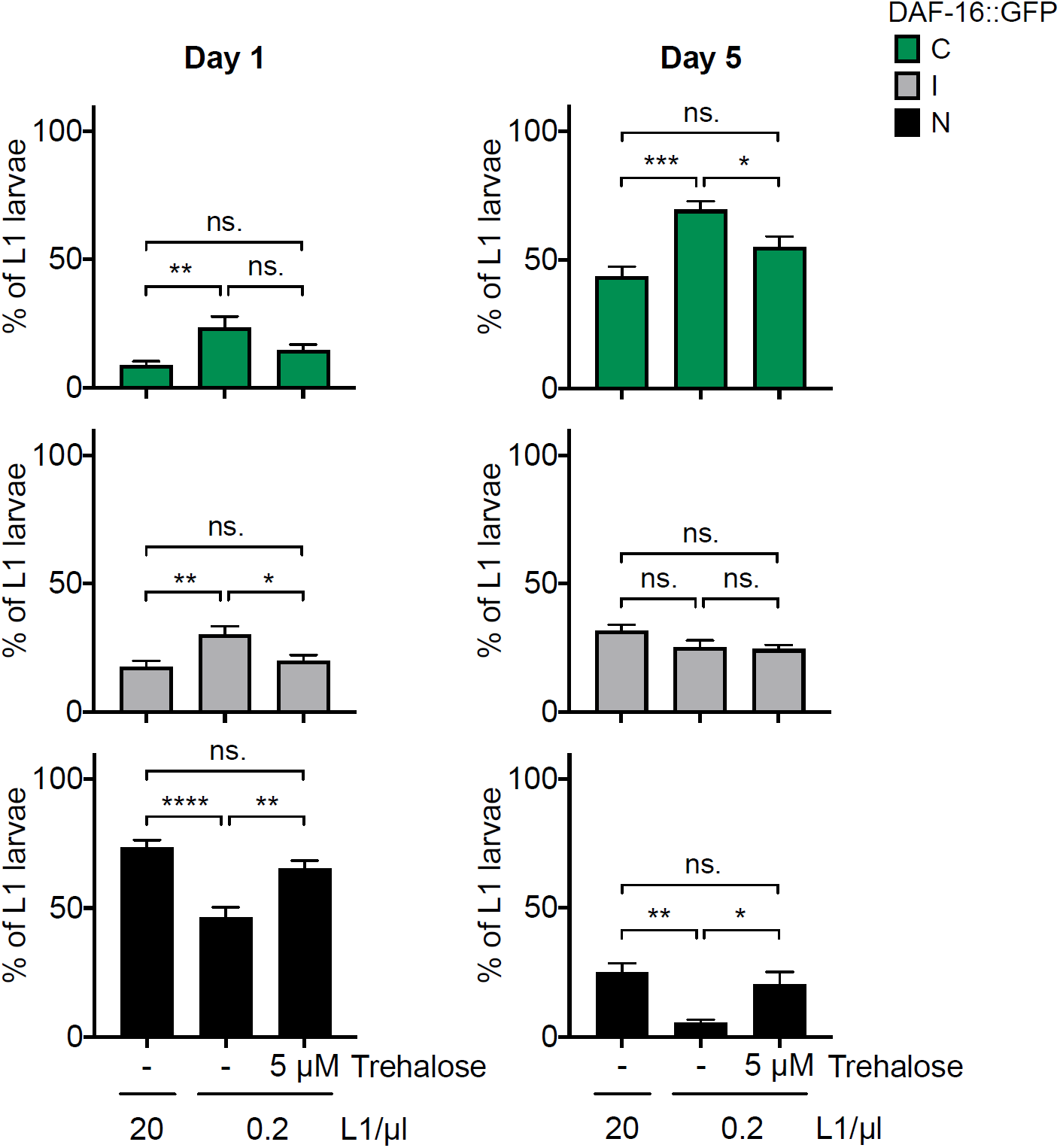
Effect on the subcellular localization of DAF-16::GFP of the addition of 5 µM of trehalose to L1 larvae arrested at 0.2 L1/µl after 1 and 5 days of starvation when compared with animals arrested at 20 and 0.2 L1/µl with no trehalose addition. Percentage of animals with cytoplasmatic localization are shown in the top panel, percentage of animals with intermediate localization are in the middle panel and nuclear localization of DAF-16 is shown in the bottom panel. The histograms show the means of eight independent replicates and the error bars represent the SEM (* p<0.05, ** p<0.01, **** p<0.0001, one-way ANOVA).

**Supplementary Figure 4.**
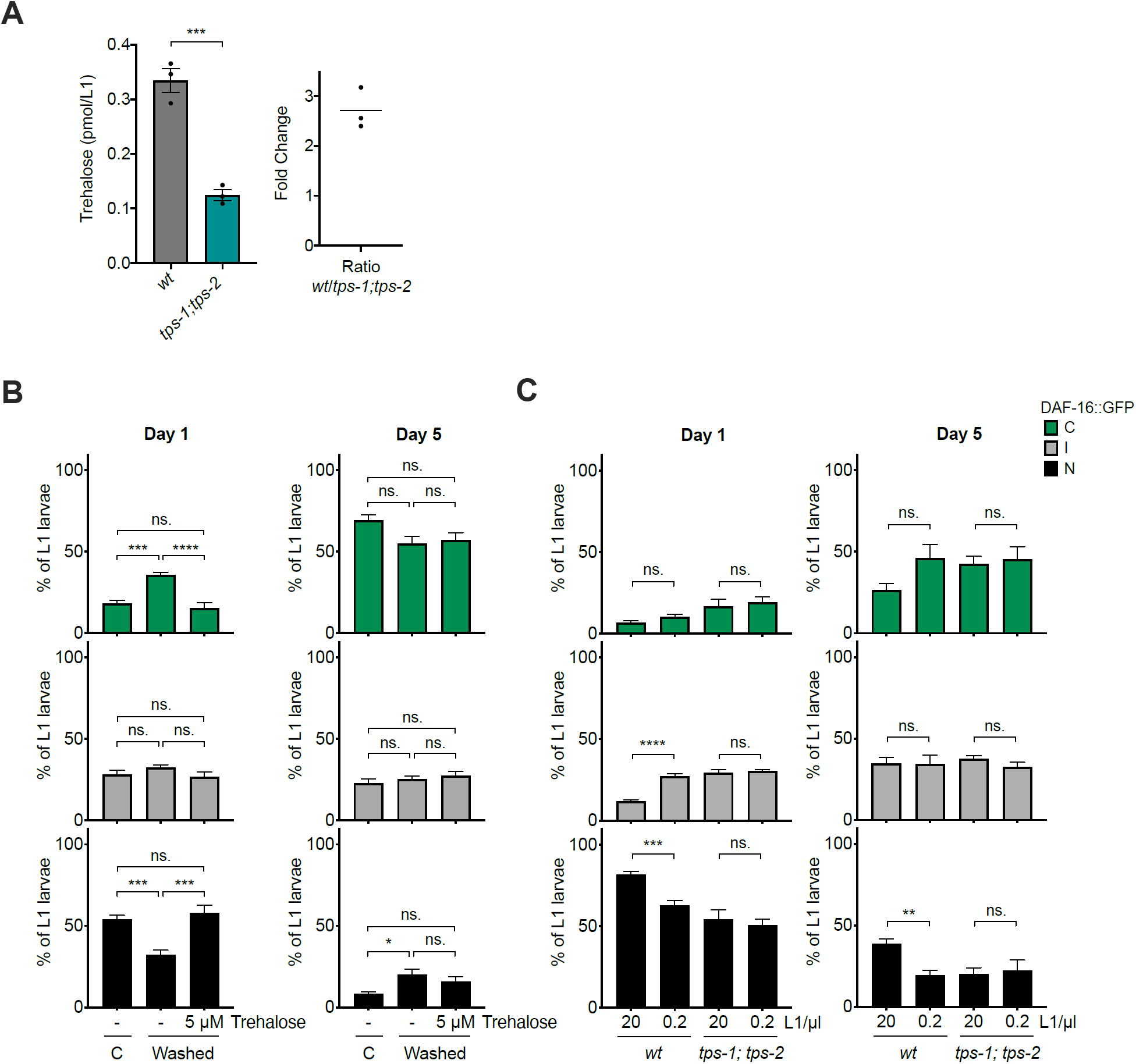
(A) In the left panel, relative trehalose content per hatched animal in the medium of wild-type and *tps-1; tps-2* strains. Black dots represent the average of each independent replicate. The bars mark the mean of the entire experiment and error bars shows the SEM. Averages of each replicate were used for statistics (*** p<0.001, Unpaired t-test). In the right panel, trehalose content ratio between wild type and *tps-1; tps-2* independently calculated for each replicate. The horizontal line indicates the mean of the three replicates. (B) Subcellular localization of DAF-16::GFP in L1 larvae arrested at 20 L1/µl at 1 and 5 days after washing the medium. Trehalose was added to one sample to a final concentration of 5 µM. In control sample the original medium was not removed and trehalose was not added. (C) Subcellular localization of DAF-16::GFP in wild-type and *tps-1; tps-2* L1 larvae arrested at 20 and 0.2 L1/µl, after 1 and 5 days of starvation. (C and D) Percentage of animals with cytoplasmatic localization are shown in the top panel, percentage of animals with intermediate localization are in the middle panel and nuclear localization of DAF-16 is shown in the bottom panel. The histograms show the means of, at least, four independent replicates and the error bars represent the SEM (* p<0.05, ** p<0.01, *** p<0.001, **** p<0.0001, one-way ANOVA in (C) and Unpaired t-test in (D)).

**Supplementary Figure 5.**
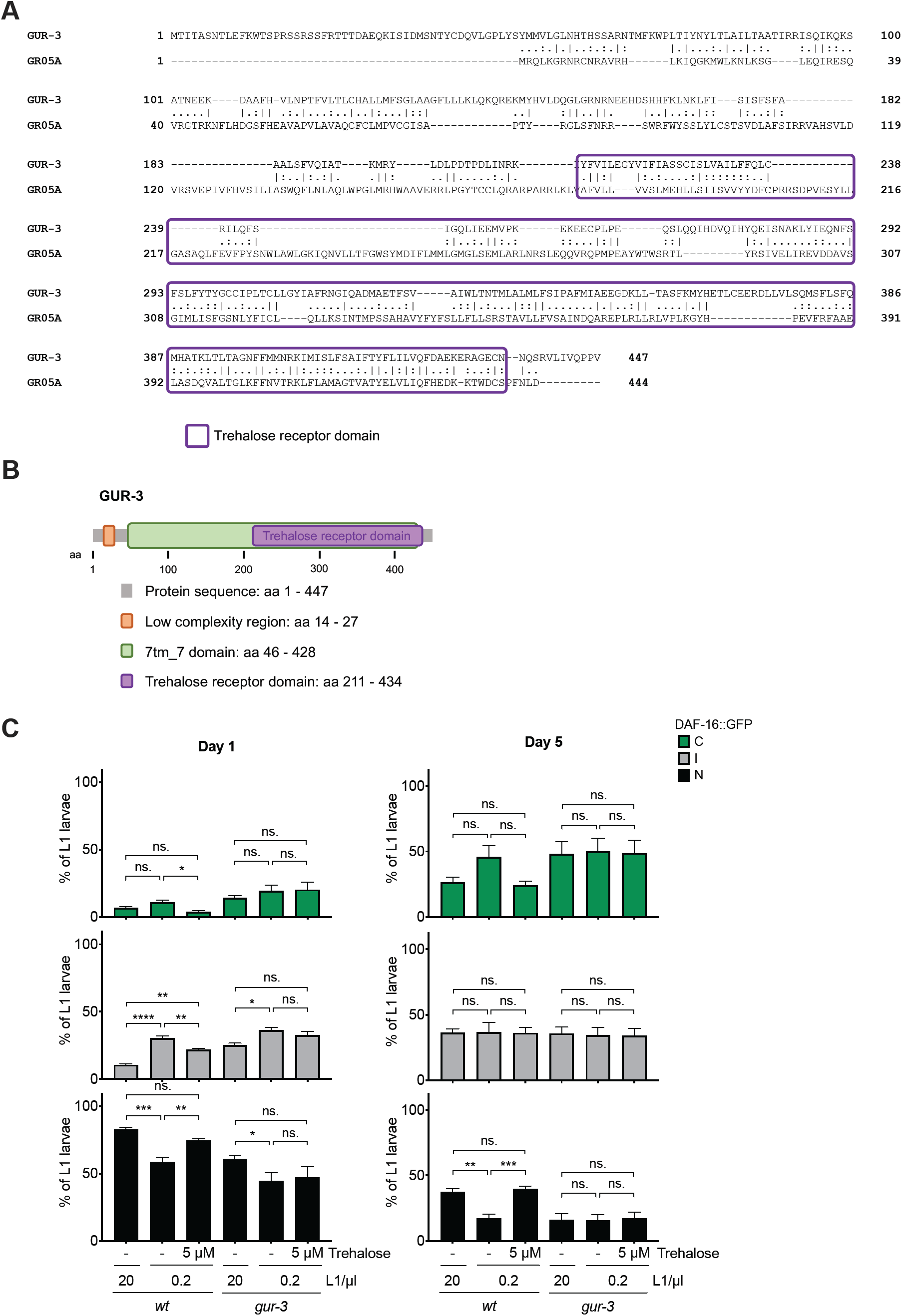
(A) Pairwise protein sequence alignment between GUR-3 and Gr5a using EMBOSS-Needle tool (https://www.ebi.ac.uk/Tools/psa/emboss_needle/). Predicted trehalose receptor domain is framed in purple. (B) Schematic representation of GUR-3 protein domains. Prediction was performed using SMART on-line tool (http://smart.embl-heidelberg.de/). (C) Subcellular localization of DAF-16::GFP in wild-type and *gur-3* L1 larvae arrested at 20 and 0.2 L1/µl after 1 and 5 days of starvation. Trehalose to a final concentration of 5 µM was added to animals at low density. Animals at high density were used as control. Percentage of animals with cytoplasmatic localization are shown in the top panel, percentage of animals with intermediate localization are in the middle panel and nuclear localization of DAF-16 is shown in the bottom panel. The histograms show the means of, at least, three independent replicates and the error bars represent the SEM (* p<0.05, ** p<0.01, *** p<0.001, **** p<0.0001, one-way ANOVA).

